# Increased Sensitivity and Signal-to-Noise Ratio in Diffusion-Weighted MRI using Multi-Echo Acquisitions

**DOI:** 10.1101/2020.03.04.962191

**Authors:** Cornelius Eichner, Michael Paquette, Toralf Mildner, Torsten Schlumm, Kamilla Pléh, Lirian Samuni, Catherine Crockford, Roman M. Wittig, Carsten Jäger, Harald E. Möller, Angela D. Friederici, Alfred Anwander

## Abstract

Post-mortem diffusion MRI (dMRI) enables acquisitions of structural imaging data with otherwise unreachable resolutions - at the expense of longer scanning times. These data are typically acquired using highly segmented image acquisition strategies, thereby resulting in an incomplete signal decay before the MRI encoding continues. Especially in dMRI, with low signal intensities and lengthy contrast encoding, such temporal inefficiency translates into reduced image quality and longer scanning times. This study introduces Multi Echo (ME) acquisitions to dMRI on a human MRI system - a time-efficient approach, which increases SNR (Signal-to-Noise Ratio) and reduces noise bias for dMRI images. The benefit of the introduced ME dMRI method was validated using numerical Monte Carlo simulations and showcased on a post-mortem brain of a wild chimpanzee. The proposed Maximum Likelihood Estimation echo combination results in an optimal SNR without detectable signal bias. The combined strategy comes at a small price in scanning time (here 30% additional) and leads to a substantial SNR increase (here up to 1.9× which is equivalent to 3.6 averages) and a general reduction of the noise bias.

## Introduction

The vast potential of dMRI in neuroscience and clinical practice led to recent developments of highly specialized human-size MRI systems with very strong diffusion gradients of up to 300mT/m (1, 2). These novel systems allow advanced dMRI acquisitions with increased signal-to-noise ratio (SNR) and, thus, higher image resolution and stronger diffusion-weightings.

Diffusion-weighted MRI (dMRI) is capable of non-invasively probing microstructure and structural connectivity of brain tissue. As a purely structural measure, dMRI can also provide insights into the organization of post-mortem brain tissue. Such post-mortem dMRI acquisitions allow extremely high image resolutions at the cost of increased scan times (3–5). Additionally, the fixation of the brain in paraformaldehyde solution produces a cross-linkage of proteins which reduces the diffusivity of water molecules in the tissue and the directional contrast (6). Hence, to achieve sufficient diffusion contrast, post-mortem diffusion-weightings need to be increased relative to in-vivo acquisitions.

Despite its versatile applications in neuroscience and clinics, dMRI acquisitions and models suffer from various shortcomings. Since the diffusion contrast is realized by direction-weighted signal attenuation, dMRI inevitably suffers from low SNR. This problem is systematically worsened for dMRI acquisitions with increased diffusion-weighting and higher image resolutions. In low SNR regimes, the typically employed magnitude dMRI signals become biased by non-Gaussian noise, thereby preventing accurate signal averaging and modeling (7–9).

Due to the time-consuming diffusion contrast encoding, dMRI data are typically acquired using Echo-Planar-Imaging (EPI). However, EPI based acquisitions are especially prone to image distortions from eddy currents or magnetic field inhomogeneities (10). A further challenge of dMRI is its inefficient signal encoding strategy, in which a rather time-consuming contrast encoding is followed by a short readout. Multiple strategies have been suggested to counteract typical issues associated with dMRI acquisitions. The signal bias of low SNR data can be overcome using phase-correction of the complex-valued dMRI dataset (8). High in-plane acceleration using parallel imaging EPI can typically reduce geometrical distortions (11–15).

For post-mortem dMRI acquisitions, these in-vivo strategies are not sufficient to achieve acceptable image quality. Captured air bubbles induce strong susceptibility differences within the post-mortem sample, making the background phase hard to estimate. In addition, post-mortem brain containers shift the tissue-air boundary from multiple centimeters to a distance of just a few millimeters. These problems aggravate EPI distortions, even when using parallel imaging acceleration.

Segmented EPI (sEPI) acquisition strategies have been proposed to mitigate these problems and to achieve almost distortion-free EPI data - even under challenging post-mortem conditions. Similar to other parallel imaging techniques, sEPI only captures small portions of the total k-space for each EPI shot. However, the missing k-space lines from segmented data undersampling are not reconstructed using additional receive coil information but rather acquired in separate shots. Hence, sEPI allows much higher acceleration factors than parallel imaging, resulting in considerably reduced image distortion. Segmented dMRI acquisitions can be achieved using interleaved segmentation along the phase encoding direction (16, 17), or by segmentation of the readout direction (18). Due to gradient slew rate limitations, interleaved segmentation along the phase encoding direction can achieve higher acceleration than readout segmented acquisitions (19).

Similarly to dMRI, fMRI recordings may also suffer from low SNR conditions, which can result in artifacts and false-positive results for BOLD activation patterns (20, 21). In this context, Multi-Echo (ME) imaging combined with acquisition acceleration strategies were shown to improve the functional sensitivity by weighted echo summation or the estimation of tissue parameters (22, 23). Despite their widespread use for functional imaging, ME acquisitions have not yet been evaluated for their applicability to diffusion-weighted MRI. In this context, short readout strategies such as sEPI can enable dense echo sampling of ME acquisitions. Hence the combination of segmented and ME acquisitions might be especially beneficial.

In this work, we present a novel dMRI sequence for postmortem acquisitions. Multiple gradient echoes of highly segmented EPI trains are acquired to achieve acquisitions with low distortions and minimal echo time. The resulting ME-dMRI signals are combined in an SNR optimal way without noise bias, using a 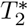 decay model and statistical modeling incorporating the underlying noise distribution. The developed acquisition strategy was employed to acquire high-resolution dMRI data of a post-mortem wild chimpanzee brain on a human-scale MRI system.

## Theory

### Description of the Sequence

The employed ME-dMRI sequence combines Stejskal-Tanner diffusion preparation (24) with repetitions of N segmented EPI readouts at different echo times TE_*n*_ (Fig. 1A). The spin-echo condition is realized at the first echo time TE_0_. Therefore, the signal in TE_*n*_ sity of *S*_0_ = *S*(TE_0_) follows a *T*_2_ decay whereas the subsequent multiple gradient-echoes decay with 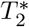 (Fig. 1B). The signal envelope of ME-dMRI data as a function of TE_*n*_ is then given by

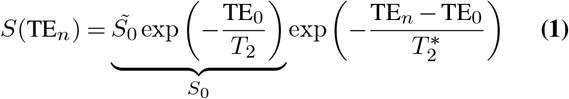

**Fig. 1.**
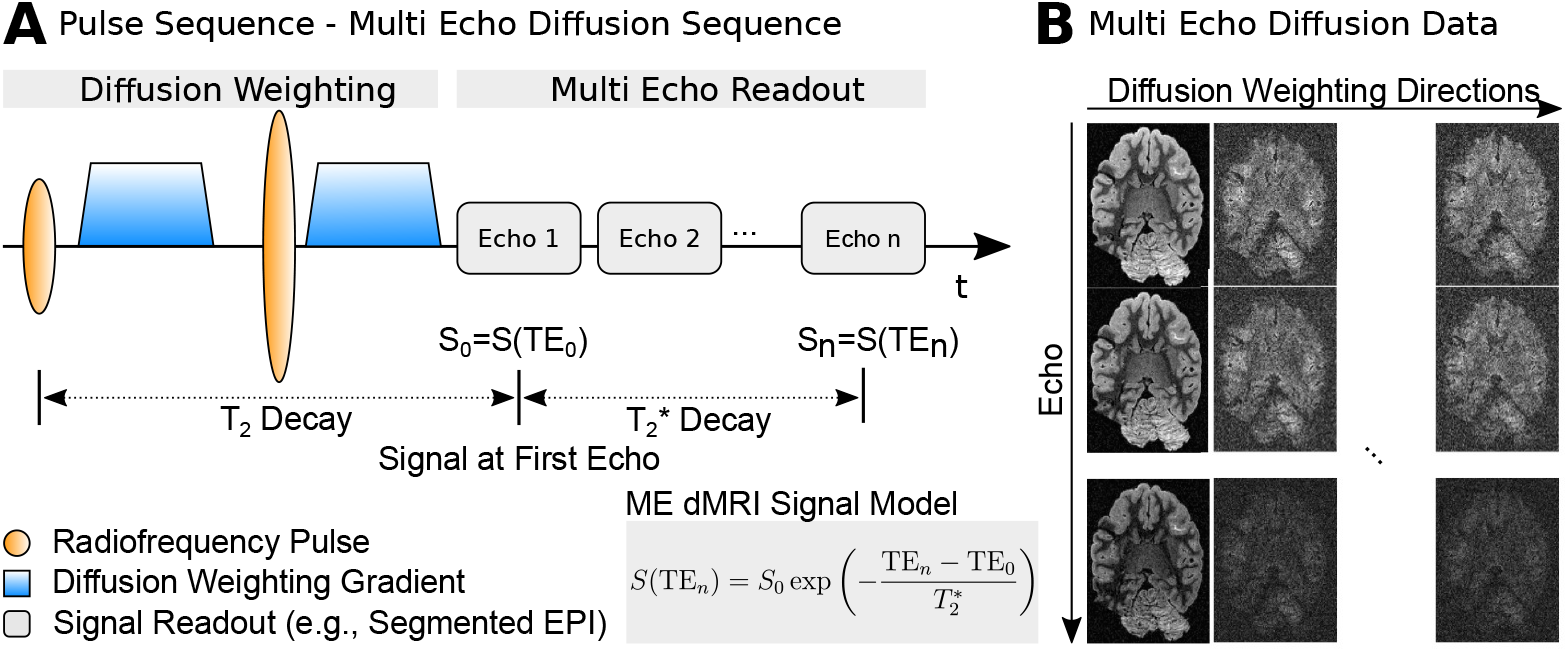
Multi-Echo dMRI Acquisitions. (**A**) ME-dMRI acquisition employs a Stejskal-Tanner dMRI sequence, followed by *N* GREs after the first echo. ME-dMRI signals are described using a mixed decay model, where the first echo follows *T*_2_ decay and the latter echoes follow 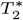 decay. (**B**) ME-dMRI data are multi-dimensional with N echo times and multiple diffusion directions.

Here, *S*_0_ describes the diffusion-weighted signal without the *T*_2_ decay component. Since this study mainly focuses on improving the SNR by acquiring multiple gradient echoes, the ME-dMRI signal decay will be considered with respect to the spin-echo signal *S*_0_:

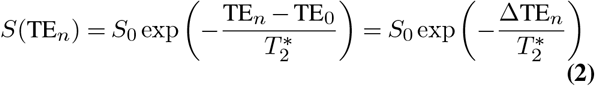

### Increasing Signal to Noise using Multi-Echo Acquisitions

ME-dMRI acquisitions provide additional data from multiple echoes which can be employed to increase the SNR. Here, we estimate the SNR gain from a ME-dMRI signal combination using the Linear Least Squares (LLS) regression formulation. We will employ two assumptions: (i) The underlying 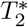 is known and will not be estimated from the ME-dMRI data. (ii) The measurement noise, *ϵ*, is assumed to follow a zero-mean Gaussian distribution with a standard deviation, *σ*. The measured signal, *M*, is described with Eq. 3:

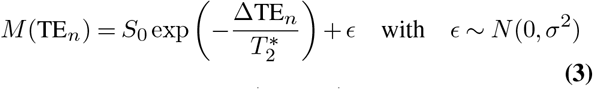

Since 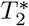 is known, 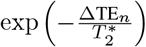, is a constant for each TE_*n*_). To estimate *S*_0_, we normalize each *M*(TE_*n*_) with this constant. It becomes apparent that the relative error of *S*_0_ estimations grows with TE. The error distribution is given by:

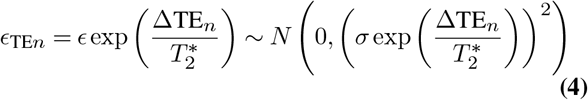

Due to the increasing error, the overall benefit of using multiple gradient echoes will depend on the parameters ΔTE, 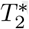, and the number of acquired echoes. If late echoes with strong error contamination are employed, the accuracy of the *S*_0_ estimation will decrease compared to early echoes. Under the assumption of Gaussian noise, *S*_0_ can be estimated as 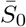 using LLS regression on multiple echoes.

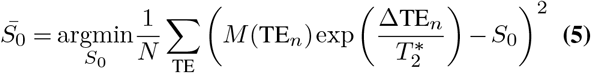

The LLS estimation of *S*_0_ is computed as the global minimum of Eq. 5.

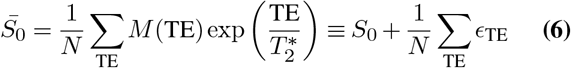

The *S*_0_ estimate from the LLS regression of ME data with Gaussian noise follows a Gaussian distribution centered around *S*_0_.

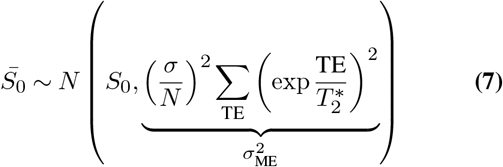

The standard deviation, *σ*_ME_, of this *S*_0_ distribution leads to an analytical expression of SNR gain from LLS ME signal estimation compared to employing only single echo data, *G*_SNR_.

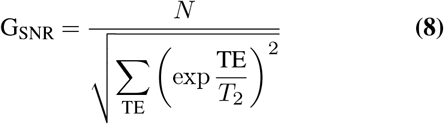

For the limiting case of infinite 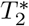, this result corresponds to the square root of N which equals the error reduction obtained by N averages. This represents the theoretical maximum SNR gain for ME-dMRI acquisitions, 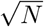. From Eq. 8 it is concluded that, for any given sampling scheme, the SNR gain is a function of tissue 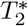. More advanced model fitting strategies such as non-linear and weighted optimization might further increase the SNR gain from using ME-dMRI data.

### Sensitivity Gain using Statistical Data Modeling

The assumption of zero-mean Gaussian noise generally does not hold for diffusion-weighted MRI, where low SNR values induce a signal bias. Therefore, for unbiased estimates of *S*_0_, it is beneficial to employ more advanced model fitting approaches, incorporating the nature of the underlying noise distribution. In contrast to LLS, the Maximum Likelihood Estimation (MLE) achieves parameter estimations of models given specific data distributions (25). By maximizing the likelihood function, MLE finds the most probable parameters of a function S to describe a given set of measurement data. MLE proves especially useful for non-Gaussian data distributions, such as dMRI data. For optimal coil combinations in complex space, dMRI signals follow a Rician data distribution characterized by the standard deviation of the underlying complex-valued noise, *σ_C_*. Using MLE, model parameters of *S*, under a Rician data distribution are computed by maximizing the logarithm of the likelihood function, *L*(26):

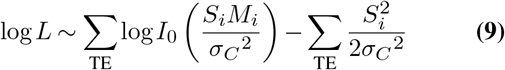

In this equation, *I*_0_ refers to the modified Bessel function of the first kind of order zero. In contrast to LLS, MLE enables unbiased estimations of model parameters by including a noise distribution model into the parameter estimation (27, 28). MLE has been shown to provide highly efficient approximations of the data by approaching the Cramér–Rao lower bound for parameter estimations (27). Due to the nonlinear weighted nature of MLE, an analytical SNR gain evaluation becomes nontrivial. We, therefore, employed numerical methods to assess the benefits of MLE parameter estimation.

## Methods

To evaluate the benefit of using Multi-Echo dMRI acquisitions, we implemented a ME-dMRI sEPI sequence and acquired data from a post-mortem chimpanzee brain. We performed numerical simulations on synthetic ME-dMRI with matching acquisition parameters to compute the SNR gain as well as potential biases from reconstructions.

### Numerical Simulations of Signal Reconstruction

Monte Carlo simulations of ME reconstructions were employed to numerically assess reconstruction accuracy for low signals and the SNR benefit from ME-dMRI, using both LLS and MLE reconstructions. To ensure comparability with the acquired ME-dMRI chimpanzee data, simulations were performed on synthetic data with a similar ME sampling scheme (i.e., numbers of echoes, echo times).

#### Evaluation of the Reconstruction Bias

Diffusion-weighted MRI acquisitions typically suffer from low SNR - especially in the main direction of diffusion attenuation, where diffusion contrast is strongest. To preserve the diffusion contrast, it is crucial that ME reconstructions retain an unbiased estimation of *S*_0_, especially for small signals. Synthetic ME-dMRI data with 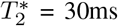 were generated for three repetitions, at five equidistantly spaced echo times from TE = 0ms to TE = 23.6ms. Signal decay curves with *S*_0_ = 1 were corrupted with complex noise. Both the real and magnitude part of the data were extracted to generate both Gaussian and Rician distributed data. Noise corrupted datasets were generated for 100 equidistantly spaced *σ*_Noise_, with *σ*_Noise, max_ = 1 to *σ*_Noise, min_ = 0.01, thereby creating decay curves at 100 different SNR levels (SNR_min_ = 1, SNR_max_ = 100). Please note that the SNR level is only valid for the first echo time, as the signal intensity decays for later echoes. LLS and MLE reconstructions of both Gaussian and Rician distributed data were performed for 1000 repetitions of each SNR level.

#### Computation of SNR Gain

The analytical assessment of SNR gain from ME-dMRI data acquisitions shows a dependency of *G*_SNR_ on the ME sampling scheme and the underlying 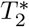 (Eq. 8). To numerically probe this relation for LLS and MLE reconstructions, we created synthetic ME data for 100 equidistantly spaced 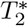 values ranging from 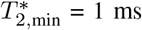 to 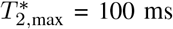. Synthetic data were generated at five equidistantly spaced echo times from TE = 0 ms to TE = 23.6 ms. Signal decay curves were corrupted with both Gaussian and Rician noise. The SNR of the noisy data was set to SNR = 5 at the first echo time. LLS and MLE reconstructions were performed for 1000 repetitions for each 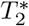 value. Reconstructions using LLS regression were performed using Eq. 6. Reconstructions using MLE regression were performed by maximizing the log-likelihood of the signal (LLS for Gaussian noise and Eq. 9 for Rician data) using the Broyden-Fletcher–Goldfarb-Shanno (BFGS) algorithm (29) as implemented in Python-SciPy (30). For each 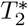 value, the SNR gain *G*_SNR_ was computed as the ratio of the standard deviation of a single echo, *σ*_Noise_, to *σ*_*S*_0__ - the standard deviation obtainable from the fitting of multiple echoes.

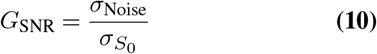

In order to employ the numerically obtained MLE SNR gains (Eq. 10) to experimental 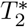 data, they were globally smoothed using a Savitzky-Golay filter and subsequently interpolated using cubic polynomials, as implemented in Python-SciPy. The SNR gain map was calculated by applying this interpolation function on the pre-calculated 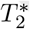 map.

### Data Acquisition

#### Specimen

MRI data were acquired from the brain of a 6-year-old juvenile wild female chimpanzee from Taï National Park, Côte d’Ivoire (31). The animal died from natural causes without human interference. The brain was extracted on-site by a veterinarian and immersion-fixed with 4% paraformaldehyde with a very short post-mortem interval of only 4 h. The performing veterinarian was specifically trained in field primate brain extractions, wearing full Personal Protective Equipment, and strictly adhering to the necropsy protocols of the field site. The procedures followed the ethical guidelines of primatological research at the Max Planck Institute for Evolutionary Anthropology, Leipzig, which were approved by the ethics committee of the Max Planck Society. The specimen was transferred to Germany strictly following CITES protocol regulations. After fixation for 6 months, the superficial blood vessels were removed, the paraformaldehyde was washed out in phosphate-buffered saline and the brain was placed for scanning in an egg-shaped acrylic container filled with perfluoropolyether (PFPE). To prevent potential leakage of PFPE during the acquisition, the container was vacuum sealed using commercially available synthetic foil packaging (Caso Design, Arnsberg, Germany).

#### MR Data Acquisition

A Stejskal-Tanner diffusion-weighted MRI sequence (24) was implemented to enable segmented EPI acquisitions of multiple echoes on a human clinical MRI system (see Fig. 1A). Diffusion-weighted MRI data of the post-mortem specimen were acquired at 3T on a MAGNETOM Skyra Connectom MRI system (Siemens Healthineers, Erlangen, Germany) using a maximum gradient strength of GMax = 300 mT/m with a slew rate of 200T/m/s and a 32 channel phased-array coil (Siemens Healthineers, Erlangen) with the following imaging parameters: 0.8mm nominal isotropic resolution, FoV = 128 × 96 mm^2^, TR = 6105ms, TE = [45.0, 50.9, 56.8, 62.7, 68.6]ms, 40 segments, Adaptive-Combine coil-combination (32), no partial Fourier, no parallel acceleration, whole-brain coverage with 80 slices. The ME-dMRI acquisition time was extended by approximately 30% compared to a Single-Echo (SE) acquisition with otherwise identical parameters. Segmented EPI echo time-shifting was employed to minimize phase-discrepancies between segments (33). Three repetitions of 60 diffusion-weighted volumes (b = 5000s/mm^2^) alongside with 7 interspersed b0 images per repetition were acquired. Due to fixation artifacts, the strong diffusion-weighting of b = 5000s/mm^2^ generated an average signal attenuation of 70%. To reduce potential impacts of magnetic field drift resulting from lengthy postmortem dMRI acquisitions, the acquisition was split into multiple 1h scans with prior frequency adjustment. To minimize impacts from short term instabilities such as vibration and tissue movement, 90 minutes of dummy scans were run prior to the actual data acquisition (5). Data from these dummy scans were discarded from all further analyses. For a statistical characterization of Rician noise of the dMRI acquisition, a noise map was recorded with identical parameters as the ME-dMRI sequence but without signal excitation (0.8 mm nominal isotropic resolution, FoV = 128 × 96mm^2^×80sl, TR = 6105ms, TE = 45.0ms, 40 segments, Adaptive-Combine coil-combination, no partial Fourier, no parallel acceleration). A 3D ME-FLASH sequence with 21 echoes was acquired for an accurate calculation of quantitative 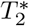 with high SNR: 0.7mm nominal isotropic resolution, FoV =112 × 100.1 × 98mm^3^, TR = 50ms, TE = 2.2 - 42.5 ms, *θ* = 32°.

### ME-dMRI Reconstruction

ME-dMRI *S*_0_ was reconstructed voxel-wise, using both LLS (Eq. 6) and MLE regression (Eq. 9). Here, all three repetitions of ME-dMRI data were jointly employed for the estimation of *S*_0_. The quantitative FLASH 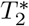 map was registered to the ME-dMRI data and included as a ground truth estimation for both LLS and MLE reconstructions. This enabled voxel-wise *S*_0_ calculation for ME-dMRI using the 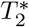 map. MLE reconstruction of Rician data requires a standard deviation estimate of the underlying complex noise, *σ*C. The whole-brain noise distribution was characterized using the acquired noise map, using a recently published method to describe multi-coil data (34). Due to the employed Adaptive-Combine coil-combination in complex space, the magnitude dMRI data approximate a Rician data distribution (35). Hence, Eq. 9 is applicable for computations of *S*_0_. MLE reconstruction was performed similarly to the preceding simulations, using BFGS optimization.

## Results

### Numerical Simulations of Signal Reconstructions

#### ME Reconstruction Accuracy

The Monte Carlo simulation results on reconstruction accuracy are summarized in Fig. 2 (top). For Gaussian distributed data, both LLS and MLE reconstruction achieve an accurate and unbiased reconstruction of the signal *S*_0_ = 1 over the full range of simulated SNR values. For Rician distributed data the LLS reconstructions of data with SNR≤10 resulted in an overestimation of *S*_0_. The simulations show that the overestimation of *S*_0_ for Rician data increases steadily with decreasing SNR values. This fact is particularly problematic for the reconstruction of dMRI data, where diffusion contrast is provided through signal attenuation. MLE reconstructions did not show overestimations of *S*_0_ at low(er) SNR values and allowed an unbiased ME reconstruction of *S*_0_ values down to SNR = 1. Monte Carlo simulations suggest that employing MLE in ME-dMRI reconstructions is favorable due to its ability to deal with low SNR values of *S*_0_.

**Fig. 2.**
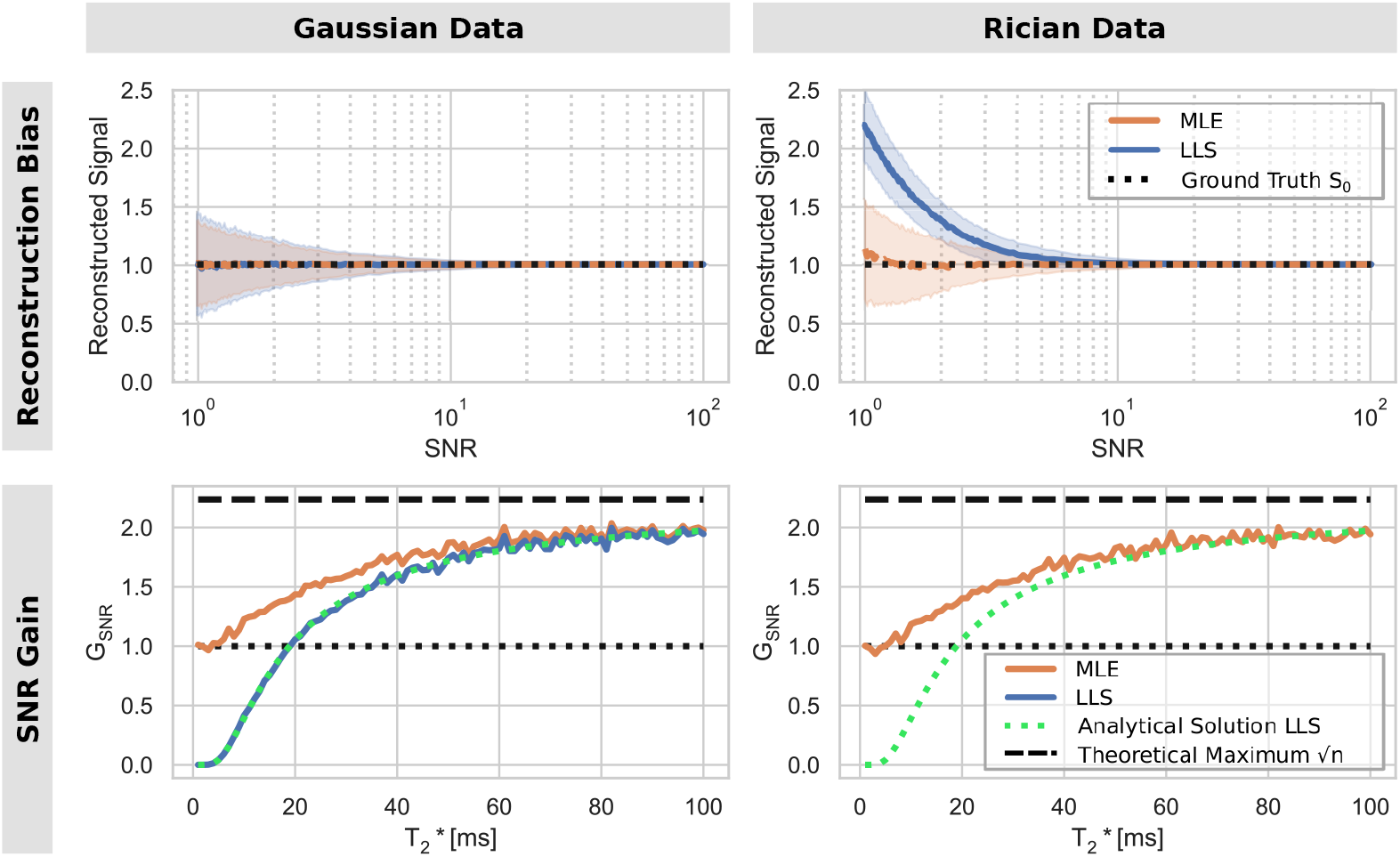
Monte Carlo Simulation Results for ME Reconstructions using MLE and LLS Regression. **Top Left:** Reconstructed *S*_0_ = 1 for 1000 repetitions for multiple Gaussian SNR levels (Log Scale). Error bars of curve show standard deviations across the simulation population. No signal bias is induced, even for very low SNR levels. **Bottom Left:** SNR Gain for LLS and MLE ME-dMRI reconstructions for Gaussian data at various 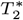 decays. Monte Carlo simulations of the LLS reconstruction display high agreement with the analytical derivation of SNR gain. MLE reconstruction achieves higher SNR gain, especially at lower 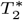 values. **Top Right:** Reconstructed *S*_0_ = 1 for 1000 repetitions for multiple Rician SNR levels. LLS reconstruction bias occurs for SNR≤10 and increases with decreasing SNR values. MLE performs an accurate estimation of *S*_0_ up to SNR = 1. **Bottom Right:** SNR Gain MLE ME-dMRI reconstructions on Rician data. MLE reconstruction achieves comparable SNR gain as for Gaussian data MLE across the entire range of 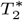.

#### SNR Gain

Figure 2 (bottom) summarizes the SNR gain, depending on the underlying 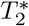 decay. Under the assumption of Gaussian noise, the results precisely follow the analytical prediction of SNR gain from Eq. 8. For growing 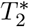 values, the SNR gain of ME-dMRI asymptotically approaches the theoretical maximum SNR gain by using 5 acquisitions (i.e., echoes), 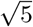. When more echoes are recorded, the theoretical maximum SNR gain will also increase. For the LLS reconstruction of ME data, the employed sampling scheme did not automatically result in an increase in SNR. Both the simulations and the analytical prediction reveal an SNR gain with LLS only for 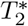 values above 20ms. This shortcoming of LLS reconstruction can be explained by the increasing error term with TEn (Eq. 4) - for very short 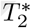 values the error of later echoes (*ϵ*_TE_) is too high to achieve an SNR gain. This can be mitigated by minimizing the echo dependent error term by means of fast and dense sampling of early echoes. MLE shows a different behavior of SNR gain across the simulated 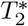 range for both the Gaussian and Rician data distributions. In both cases, SNR loss was not observed - even for short 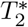 values. Instead, the SNR gain converges to *G*_SNR_ = 1 for small 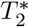 values, where neither SNR gain nor loss will occur. Furthermore, MLE reconstruction achieves a greater SNR gain than LLS over the entire range of simulated 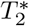 values. MLE on Gaussian data reduces to a least-squares problem, which is similar to the LLS algorithm. Therefore, the SNR gain compared to the LLS computation from Equation 6 might be attributed to the employed nonlinear BFGS optimization algorithm. The key advantage of MLE optimization is the ability to achieve SNR gains > 1 over the whole 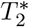 range, especially for Rician distributed data. In summary, MLE reconstruction enables accurate and unbiased reconstructions of *S*_0_ with high SNR. Monte Carlo simulations suggest that MLE is an optimal reconstruction method for reconstructing ME-dMRI data.

#### ME-dMRI Reconstruction

Figure 3A compares reconstruction results from Single-Echo and Multi-Echo dMRI acquisitions. The raw data quality of the first echo is shown in the top row. The SNR gain of the acquired ME-dMRI data (three repetitions, five echoes) becomes visible in comparison with the SE-dMRI data (three repetitions, single echo). When comparing ME-dMRI reconstructions, it is apparent that the diffusion attenuation contrast is considerably more pronounced for MLE reconstructions. Therefore, the ME-dMRI reconstruction results support the previous simulation results by demonstrating that MLE regression prevents signal bias for small *S*_0_ values. This becomes especially visible in the diffusion-weighted data, where the diffusion induced signal attenuation is much more pronounced. Figure 3B shows the signal intensity distribution for MLE and LLS ME-dMRI reconstructions. Noise-induced bias becomes visible when comparing the signal distributions across the whole-brain volume for both diffusion-weighted and non-diffusion-weighted volumes. Here, LLS reconstructions show a tendency towards increased image intensities - i.e., a weakening of the diffusion contrast compared to MLE.

**Fig. 3.**
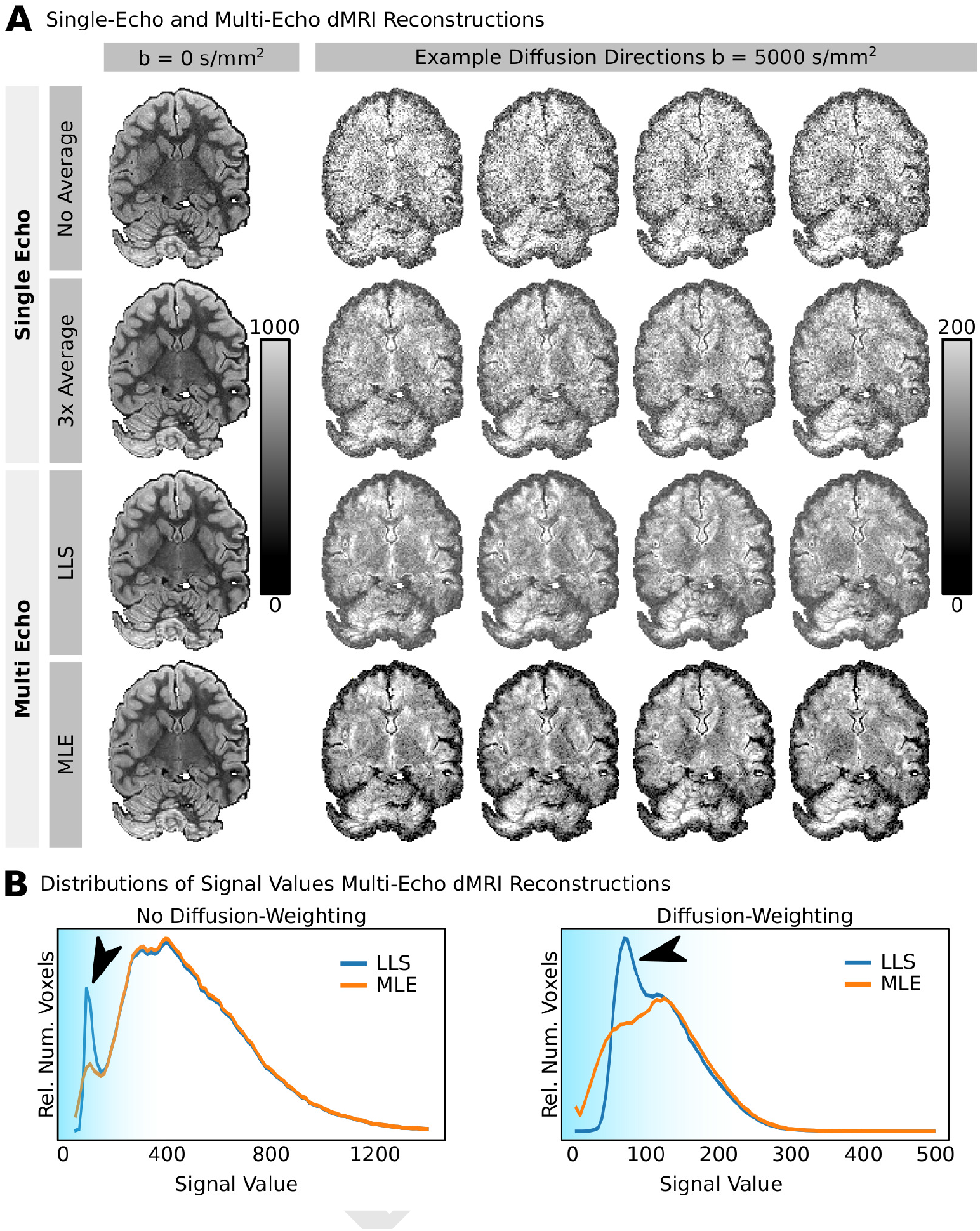
Exemplary Single-Echo and Multi Echo Reconstructions for B0 and Exemplary Diffusion Directions. (**A**) Single-Echo: The non-averaged Single-Echo diffusion-weighted data display very poor SNR (top row). Threefold averaging of dMRI data introduces improvements of image SNR scaled by 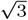 (second row). Multi Echo: Multi Echo dMRI acquisitions displays clearly visible SNR benefits. When comparing LLS (row three) and MLE reconstructions (row four), the benefits of noise informed MLE reconstructions become evident. MLE reconstructions display a much stronger diffusion contrast due to a better reconstruction of diffusion signal attenuation. (**B**) Distributions of reconstructed ME-dMRI data show differences, only for low signal intensities in both dMRI data without (left) and with diffusion-weighting (right).

#### SNR Gain Map

Figure 4 displays the whole-brain 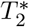 map as well the associated SNR gain from using ME-dMRI alongside with MLE reconstruction. Given a specific ME sampling scheme the voxel-specific SNR is defined by the underlying 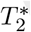 map (Fig. 4A). The quantitative 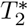 measurements show a distribution, ranging from 20ms to 80ms across the brain (Fig. 4B). The numerically generated SNR gain for this 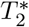 distribution using MLE is summarized in Fig. 4C. The SNR gain from a single repetition of ME-dMRI shows a tissue-specific SNR gain of 1.6 (WM, equivalent to 2.6 averages) and 1.9 (GM, equivalent to 3.6 averages). From the *G*_SNR_ histogram in Fig. 4D, it is apparent that different tissue types generate distinct SNR gains. Inclusions of air bubbles can reduce 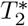 within a small radius as a result of susceptibility differences. In such areas, the Multi-Echo combination does not allow significant SNR gain, as 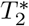 is too short. Please note that these results are shown for a single Multi-Echo combination. Experimentally, a total of three repetitions were recorded, increasing the final SNR by an additional factor of 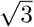.

**Fig. 4.**
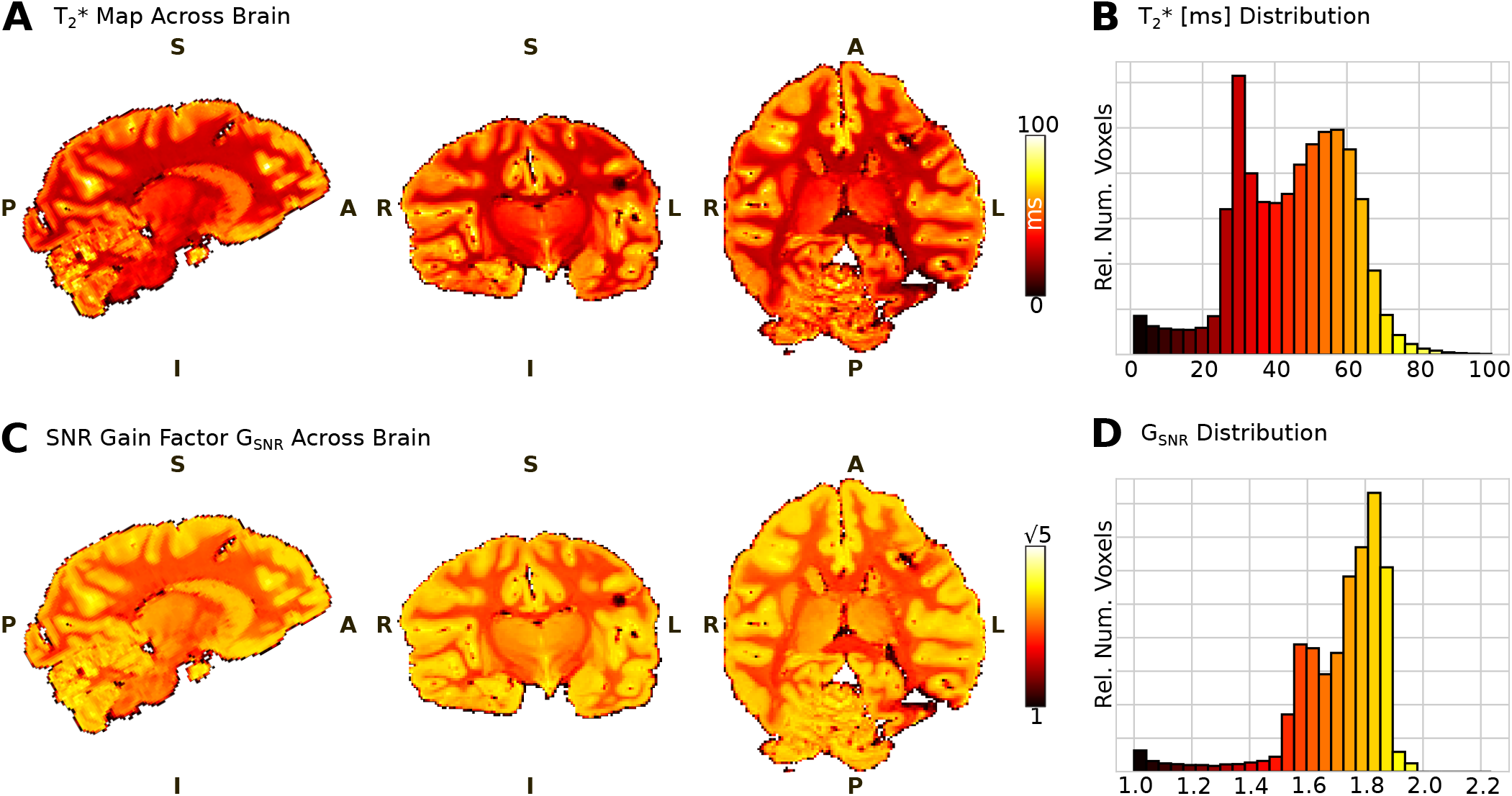
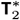-dependent SNR Gain of ME-dMRI Acquisitions. (**A**) Whole-brain 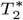 map in orthographic view. (**B**) The 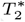 histogram displays a bimodal 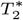 distribution across the brain with WM 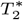 30ms and GM 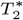 60ms. Voxels with very low 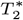 values (< 25ms) are likely caused by residual air bubbles. (**C**) Whole-brain map of 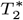 dependent SNR gain. GSNR of ME-dMRI is strongest in areas of long 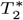, such as in GM. The SNR gain map is windowed to the theoretical SNR gain of 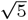 from acquired 5 acquisitions (echoes). (**D**) Histograms of SNR gain from one repetition of ME-dMRI show a tissue-dependent SNR gain of 1.6 (WM, equivalent to 2.6 averages) and 1.9 (GM, equivalent to 3.6 averages). The whole-brain SNR gain increases if shorter readout trains enable higher density sampling of the first echoes.

## Discussion

In dMRI acquisitions, accurate and unbiased measurements of signal attenuation are key to reconstruct fiber orientations or diffusion models with sufficient accuracy. Diffusion-weighted MRI measurements are based on selective signal attenuation and consequently suffer from intrinsically low SNR. In advanced dMRI, spatial resolution and diffusion-weighting are continuously increased (36–38). Such measurements more accurately capture the structure of the brain and allow better characterizations of underlying tissue microstructure. However, low SNR regimes are a limiting factor in such advanced dMRI acquisitions, thereby dampening the diffusion contrast and accurate model estimations from such measurements.

Here we present a novel strategy to increase SNR and reduce the signal bias from non-Gaussian noise for advanced dMRI acquisitions with low SNR. In this context, we developed a diffusion-weighted sEPI sequence capable of recording Multi-Echo signals following each diffusion preparation period. The Multi-Echo data are reconstructed using a signal relaxation model and a quantitative 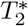 map. The additionally required 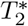 map can be recorded within a few minutes - a time effort disproportionate to typically time-consuming post-mortem dMRI measurements.

To ensure accurate and optimal reconstructions of ME-dMRI data, we performed numerical Monte-Carlo simulations of different reconstruction algorithms and data types across various signal parameters. The simulations demonstrated optimal SNR gain and minimal signal bias using Maximum Likelihood Estimation reconstruction incorporating the Rician distribution of the data. Both simulations and analytical evaluation of the optimization problem showed an SNR gain dependency on both underlying 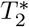 and the Multi-Echo sampling density. In our case, the SNR gain of white matter was 1.6 - which is equivalent to 2.6 averages. For grey matter, the SNR gain was 1.9 - equivalent to 3.6 averages. The SNR gain of up to 3.6 averages comes at the cost of an only slightly extended measurement time of approximately 30%.

The second advantage of ME reconstructions using MLE is the low signal bias. Since MLE can incorporate the underlying noise distribution into the S0 computation, the reconstructed signal remains unbiased, even for very low SNR levels of SNR 1. An LLS reconstruction of the same data led to signal bias already present at SNR = 10. Therefore, MLE allows an unbiased reconstruction of the ME-dMRI data of signals with attenuations of almost an additional order of magnitude. This is particularly important in diffusion imaging, where low signals must be calculated with maximum precision.

To put our proposed method to direct use, we recorded postmortem dMRI data of a wild chimpanzee brain with high resolution and strong diffusion-weighting. The reconstruction of the acquired ME-dMRI data resulted in high-quality results with a noticeable SNR gain. In addition, the ME-dMRI reconstructions are in agreement with the simulations such that the desired diffusion contrast strongly increased when MLE reconstruction was used.

The implementation of this sequence on a human-scale MRI system capable of a maximum gradient strength of 300mT/m allowed the acquisition of dMRI data with high diffusion-weightings to study the connectivity of whole brains up to human size. Thereby, sufficient directional diffusion contrast was achieved - also in post-mortem tissue, which requires stronger b-values to achieve similar contrast compared to in-vivo tissue.

The high spatial resolution of the acquired dMRI data provides the basis for a more accurate reconstruction of the white matter tracts compared to typical in-vivo scans. This reduces the potential pitfalls of diffusion MRI tractography (39) and allows reconstructing the endpoints of the fiber pathways with greater precision. This is of particular relevance in the evolutionary context of the current study.

The strong in-plane acceleration of the sEPI acquisition allows minimizing image distortions which are caused by susceptibility differences. This enables more precise measurements of anatomical structures and is most relevant for postmortem acquisitions, which may include captured air bubbles and strong susceptibility contrasts at the edges of the brain container.

The discussed ME approach is not limited to post-mortem dMRI sEPI acquisitions. The concept of ME-dMRI could also be used to increase the SNR and reduce signal bias in other settings. Segmented EPI dMRI acquisitions might also be acquired in-vivo, utilizing phase-navigators (18, 40) or advanced reconstruction mechanisms (41, 42). ME-dMRI is particularly advantageous for accelerated sequences with a very short EPI readout time, where the signal of the multiples echos is still sufficient considering the 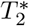 decay. Therefore, with strong in-plane acceleration, in-vivo applications of ME-dMRI data might also be feasible.

ME-dMRI acquisition schemes are not exclusively applicable by using sEPI. Instead, they can also be extended to other dMRI acquisition strategies such as steady-state free precession (SSFP) dMRI (3) to increase the SNR and reduce noise bias of the data. Due to the non-EPI single line readout of SSFP dMRI sequences, the SNR increase could be even larger due to a denser sampling of early echoes.

Furthermore, the concept of noise informed decay modeling is applicable to all MRI modalities where non-Gaussian noise can induce signal biases. Noise informed MLE fitting can be beneficial for the quantitative mapping of magnetic tissue properties. For example, noise informed MLE reconstruction could be advantageous for other quantitative MRI strategies using Multi-Echo acquisitions such as ME-FLASH (43) or ME-MP2RAGE (44). Multi-Echo reconstructions are commonly performed using LLS (45, 46). Especially at high resolutions or for post-mortem acquisitions, signal bias effects may be circumvented using advanced regression strategies such as MLE. We expect advanced fitting strategies to also be beneficial to diffusion relaxometry, where signal attenuation is generated not only from diffusion-weighting but also from additional mechanisms such as *T*_1_ relaxation and *T*_2_ relaxation (47–49).

## Conclusion

Here we present Multi-Echo dMRI - a new concept to both increase image SNR and simultaneously reduce signal-biases of noise-corrupted dMRI data for post-mortem acquisitions. Diffusion MRI acquisitions typically suffer from a low temporal encoding efficiency, where a time-consuming contrast encoding is followed by a rather rapid acquisition of only one signal. Through the rapid sequential recording of multiple echoes, this new acquisition and reconstruction strategy makes better use of the diffusion-weighted signal and results in a more time-efficient contrast encoding. The presented ME-dMRI technique might also be beneficial to in-vivo dMRI acquisitions as well as quantitative relaxometry.

## ACKNOWLEDGEMENTS

CE and MP are supported by the SPP 2041 ‘Computational Connectomics’ of the German Research Foundation, DFG. MP is supported by a scholarship (PDF-502732-2017) from the Natural Sciences and Engineering Research Council of Canada (NSERC). The Max Planck Society provides core funding for the Taï Chimpanzee Project since 1997. We are grateful to the Ministère de l’Enseignement Supérieur et de la Recherche Scientifique and the Ministère de Eaux et Forêts of Côte d’Ivoire and the Office Ivorien des Parcs et Réserves for permissions to conduct the research; also to the Centre Suisse de Recherches Scientifiques for their long-term support and the staff members of the Taï Chimpanzee Project for their continuous work in the field.

